# Integrated Spatial and Single-Nuclei Transcriptomic Analysis of Long Non-Coding RNAs in Alzheimer’s Disease

**DOI:** 10.1101/2024.10.27.620550

**Authors:** Bernard Ng, Denis R. Avey, Faraz Sultan, Katia de Paiva Lopes, Masashi Fujita, Devin Saunders, Ricardo A. Vialle, Himanshu Vyas, Nicola A. Kearns, Shinya Tasaki, Artemis Iatrou, Sashini De Tissera, Amiko Krisa Lagrimas, Tien-Hao Chang, Jishu Xu, Chunjiang Yu, Vilas Menon, Chris Gaiteri, Philip L. De Jager, David A. Bennett, Yanling Wang

**Affiliations:** Rush Alzheimer’s Disease Center, Rush University Medical Center, Chicago, IL, 60612, USA; Center for Translational and Computational Neuroimmunology, Department of Neurology, Columbia University Irving Medical Center, New York, NY, USA; Current affiliation: Department of Psychiatry, McLean Hospital, Harvard Medical School, Belmont, MA, 02478, USA; Department of Psychiatry, Upstate Medical University, Syracuse, NY, 13210, USA; Department of Neurological Sciences, Rush University Medical Center, Chicago, IL, 60612, USA

**Keywords:** long non-coding RNA, spatial transcriptomics, snRNA-seq, Alzheimer’s disease, OIP5-AS1, microglia

## Abstract

**Background:** Long non-coding RNAs (lncRNAs) are critical regulators of physiological and pathological processes, with their dysregulation increasingly implicated in aging and Alzheimer’s disease (AD). To investigate the spatial and cellular distribution of lncRNAs in the aging brain, we leveraged published spatial transcriptomics (ST), single-nucleus RNA sequencing (snRNA-seq), and bulk RNA-seq datasets from the dorsolateral prefrontal cortex (DLPFC) of ROSMAP participants with and without pathological AD.

**Results:** LncRNAs exhibited greater subregion-specific expression than mRNAs, with enrichment in antisense and lincRNA biotypes. Subregion-enriched lncRNAs were generally not cell-type specific, and vice versa. Differential expression analysis of ST data identified AD-associated lncRNAs with distinct spatial patterns and moderate overlap with differentially expressed (DE) lncRNAs from bulk RNA-seq. Gene set enrichment revealed their involvement in chromatin remodeling, epigenetic regulation, and RNA metabolism. We also identified AD DE lncRNAs across major brain cell types using snRNA-seq but overlap with ST DE lncRNAs was limited. Among previously reported lncRNAs, *OIP5-AS1* was consistently upregulated in AD in all cortical subregions. Antisense oligonucleotide (ASO) knockdown of *OIP5-AS1* in iPSC-derived microglia led to upregulation of pro-inflammatory genes and downregulation of DNA replication and repair pathways. Immunoassays confirmed increased secretion of pro-inflammatory cytokines. The knockdown expression pattern was enriched for microglia-specific AD DE genes and microglia states.

**Conclusions:** This study provides a spatial and cellular map of lncRNAs in the aging human cortex and identifies subregion-and cell-type-enriched DE lncRNAs in AD. Our findings implicate *OIP5-AS1* in microglial activation, suggesting its potential contribution to AD pathogenesis.

## Introduction

One of the most remarkable discoveries of the 21st century is that a large portion of the genome is transcribed into RNAs with little or no protein-coding potential (1,2). While approximately 90% of the human genome is transcribed, only about 2% is annotated as protein-coding genes. This paradigm-shifting discovery has spurred extensive research into the regulatory roles of non-coding RNAs in gene expression (2,3), redefining our understanding of the genome and challenging the traditional view that only protein-coding genes are functionally relevant.

Among non-coding RNAs, long non-coding RNAs (lncRNAs) represent a prominent class of transcripts longer than 200 nucleotides, predominantly transcribed by RNA Polymerase II. LncRNAs regulate gene expression through diverse mechanisms, including epigenetic modification, chromatin remodeling, and post-transcriptional processes (4,5). Compared to coding RNAs, lncRNAs exhibit greater tissue and cell-type specificity (6,7) and are particularly abundant in the human brain, where they play critical roles in neuronal differentiation, synaptogenesis, and neuroplasticity (1,8). Dysregulated lncRNA expression has been implicated in numerous neurological disorders, including Alzheimer’s disease (AD). In AD, aberrant lncRNA expression has been linked to key pathological processes, such as amyloid-beta metabolism, tau pathology, neuroinflammation, and neuronal death (9–13). However, the spatial and cellular contexts in which lncRNAs might influence AD-related processes remain largely unexplored.

In this study, we leveraged our published spatial transcriptomics (ST) data (14) as well as published single-nucleus RNA sequencing (snRNA-seq) (15) and bulk RNA-seq (16) datasets from the dorsolateral prefrontal cortex (DLPFC) of ROSMAP participants to investigate the spatial and cellular expression of lncRNAs in the aging brain. We identified subregion-specific lncRNAs and assessed their overlap with cell-type-specific lncRNAs. Additionally, we mapped lncRNAs and mRNAs that are differentially expressed (DE) in AD across distinct subregions and compared the ST findings with results from bulk RNA-seq and snRNA-seq analyses. Finally, we functionally tested an AD-associated lncRNA, *OIP5-AS1*, in iPSC-derived microglia using antisense oligonucleotide knockdown and revealed its potential roles in AD pathogenesis.

## Results

### Spatial expression of lncRNAs in aged human brains

Using our recently published ST data (14), we analyzed lncRNA expression from 78 DLPFC sections of 21 female individuals (AD = 13; no cognitive impairment (NCI) = 8). In total, we detected 7,634 lncRNAs and 16,769 protein-coding mRNAs across 8 spatial spot clusters corresponding to subregions: layer 1 (L1), layers 2–3 (L2–3), layers 3–5 (L3–5), layer 5 (L5), layer 6 (L6), white matter (WM), meninges, and blood vessels (Fig. 1A). Consistent with prior studies (6,7), lncRNAs exhibited lower expression levels than mRNAs (Fig. 1B-C). For downstream analysis, we retained 1,578 lncRNAs and 13,808 mRNAs with adequate expression (counts per million [CPM] >1 in >80% of brain sections). Among these lncRNAs, the most prevalent biotypes were long intergenic non-coding RNAs (lincRNAs) and antisense transcripts (Fig. 1D). Notably, lncRNAs demonstrated greater subregion specificity, with only 39.9% of lncRNAs detected across all eight subregions, compared to 89.3% of mRNAs (Fig. 1E).

**Figure 1.**
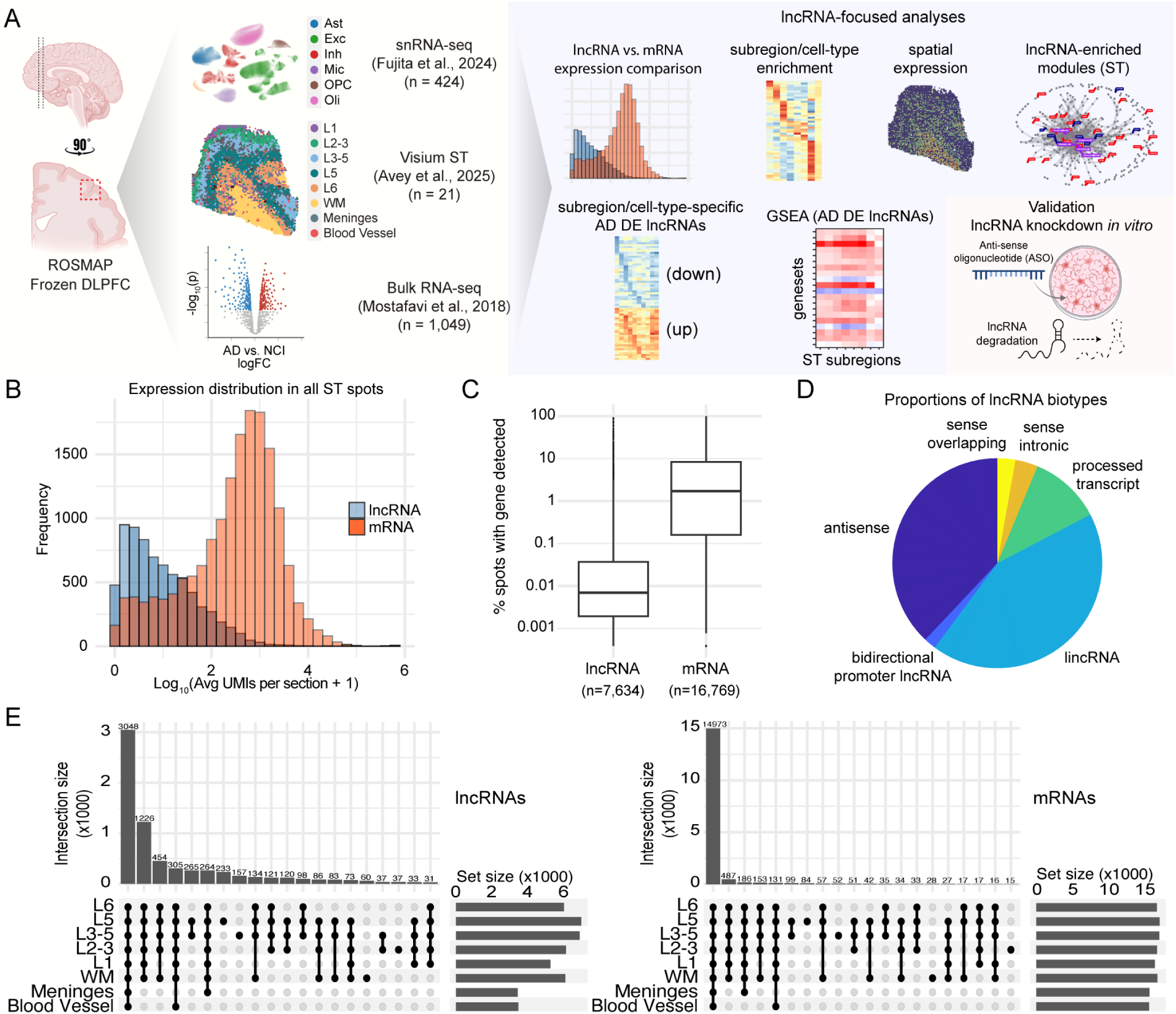
Spatial expression of lncRNAs. **(A)** Study overview. **(B)** Comparison of the expression distributions of lncRNAs and protein-coding genes (mRNAs). **(C)** Comparison of percentage of ST spots with non-zero expression for lncRNAs and mRNAs. **(D)** Proportion of detected lncRNAs (aggregated across subregions) belonging to each biotype. **(E)** Upset plots showing the numbers of lncRNAs (left) and mRNAs (right) detected in the top 20 most common intersections of subregions.

As a data quality check, we investigated the molecular systems within each subregion by generating a pseudo-bulk matrix for each subregion and performing co-expression network analysis using the Speakeasy algorithm(17) (Fig. S1). This analysis generated 193 modules, each containing at least 30 genes (Table S2). Module preservation analysis revealed that cortical layer modules were more preserved than those derived from WM, blood vessels, and meninges (Fig. S1B; Table S3). Within each subregion, 15–60% of the modules were lncRNA-enriched (Fig. S1) with many enriched for oxidative phosphorylation, synaptic signaling, immune response, cell cycle regulation, and ncRNA processing (Table S4), in line with the functional roles of ncRNA.

### Comparison of subregion-and cell-type-specific lncRNA expression

By contrasting the expression of spatial spots in one subregion against all others in our ST data, we identified 79 subregion-specific lncRNAs (Table S1). To investigate whether subregion-specific lncRNAs also exhibit cell-type specificity, we cross-referenced a published snRNA-seq dataset (15) derived from the DLPFC of 424 ROSMAP participants(15). We observed limited overlap between subregion-specific and cell-type-specific lncRNAs (Fig. 2A; Table S1), but some exceptions emerged. For example, *AL031056.1* was enriched in L1 and astrocytes (Fig. 2C), *LINC00507* in L2–3 and excitatory neurons (Fig. 2B-C), *AC021613.1* in L6 and excitatory neurons (Fig. 2B-C), and *AC092958.1*, *TYMSOS*, and *LINC00844* in WM and oligodendrocytes (Fig. 2B-C). However, most subregion-specific lncRNAs were expressed in all major cell types (Fig. 2C).

**Figure 2.**
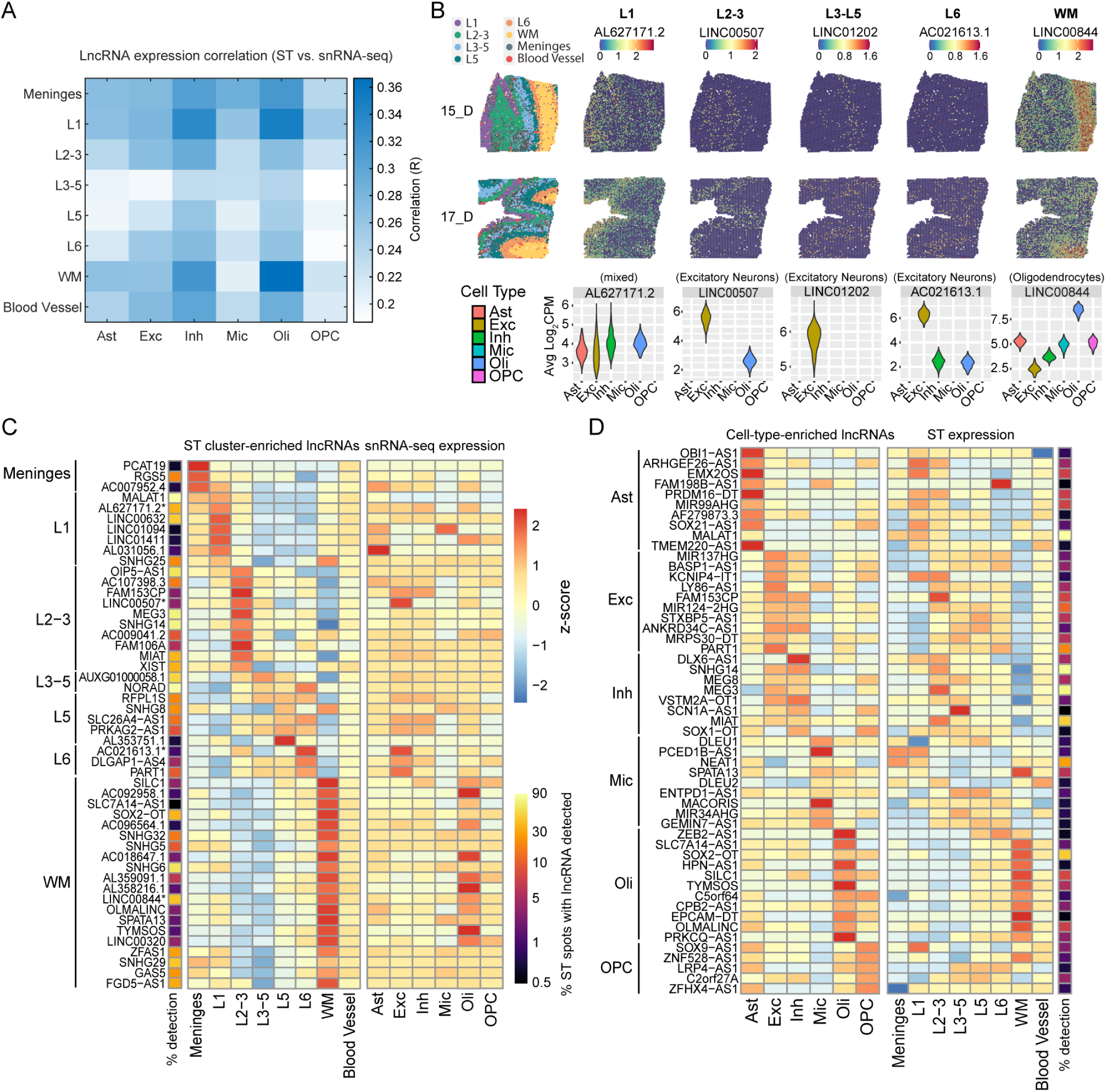
Enrichment of lncRNA expression by spatial subregion or cell type. **(A)** Correlation heatmap of ST vs. snRNA-seq expression among all co-detected lncRNAs. **(B)** UMAP and expression feature plots for two representative sections, showing 5 examples of lncRNAs with enriched expression in specific cortical layers or white matter (WM). Violin plots below show expression distributions among 6 major brain cell types. **(C)** Heatmap showing scaled expression (z-score) of the top 50 subregion-specific lncRNAs (left), and their cell-type-specific expression (right). The first column shows the percentage of spots with non-zero expression for each lncRNA. (*asterisks indicate the lncRNAs shown in panel B). **(D)** Scaled expression by cell type (left) and ST subregion (right) for the top 10 cell-type-specific lncRNAs from each of the 6 major brain cell types.

We next examined whether cell-type-specific lncRNAs exhibited subregion specificity. Although general trends emerged, such as astrocyte-specific lncRNAs being more abundant in L1, excitatory/inhibitory neuron-specific lncRNAs in cortical layers, and oligodendrocyte-specific lncRNAs in WM, many cell-type-specific lncRNAs were expressed across multiple subregions (Fig. 2D; Table S1).

### AD DE lncRNAs revealed by ST

To identify AD DE lncRNAs in our ST data, we aggregated spot-level expression values within each brain section to generate a pseudo-bulk matrix for each subregion. We then normalized these estimates using TMM-voom (18), and applied linear mixed models (LMM) to contrast AD against NCI. We accounted for age, RNA Integrity Number (RIN), postmortem interval (PMI), and batch as fixed effects, and within-subject correlation as a random effect. We declared differentially expressed genes at an *ɑ* of 0.05 with false discovery rate (FDR) correction across genes and subregions.

We identified 275 AD DE lncRNAs (Table S5). A greater proportion of these lncRNAs exhibited negative associations with AD compared to mRNAs (Fig. 3A-B) and biotype analysis revealed enrichment of bidirectional promoter lncRNAs in L3–5 and lincRNAs in L6 and WM (Fig. 3C). Next, we compared our ST-derived AD DE lncRNAs with those identified from bulk RNA-seq data of the DLPFC in 1,049 individuals (16) (Table S5). Although the overlap of AD DE genes between ST and bulk RNA-seq was modest (Fig. 3D), the set of t-values across their common lncRNAs exhibited moderate correlations between the two modalities (R = 0.3; Fig. 3D). Concordant findings included several downregulated lncRNAs, such as *LY86-AS1*, *FAM106A*, *LINC01007*, and *MIR7-3HG*, as well as upregulated lncRNAs, including *NEAT1*, *LINC00632*, *FGD5-AS1*, and *LINC01094* (Fig. 3D).

**Figure 3.**
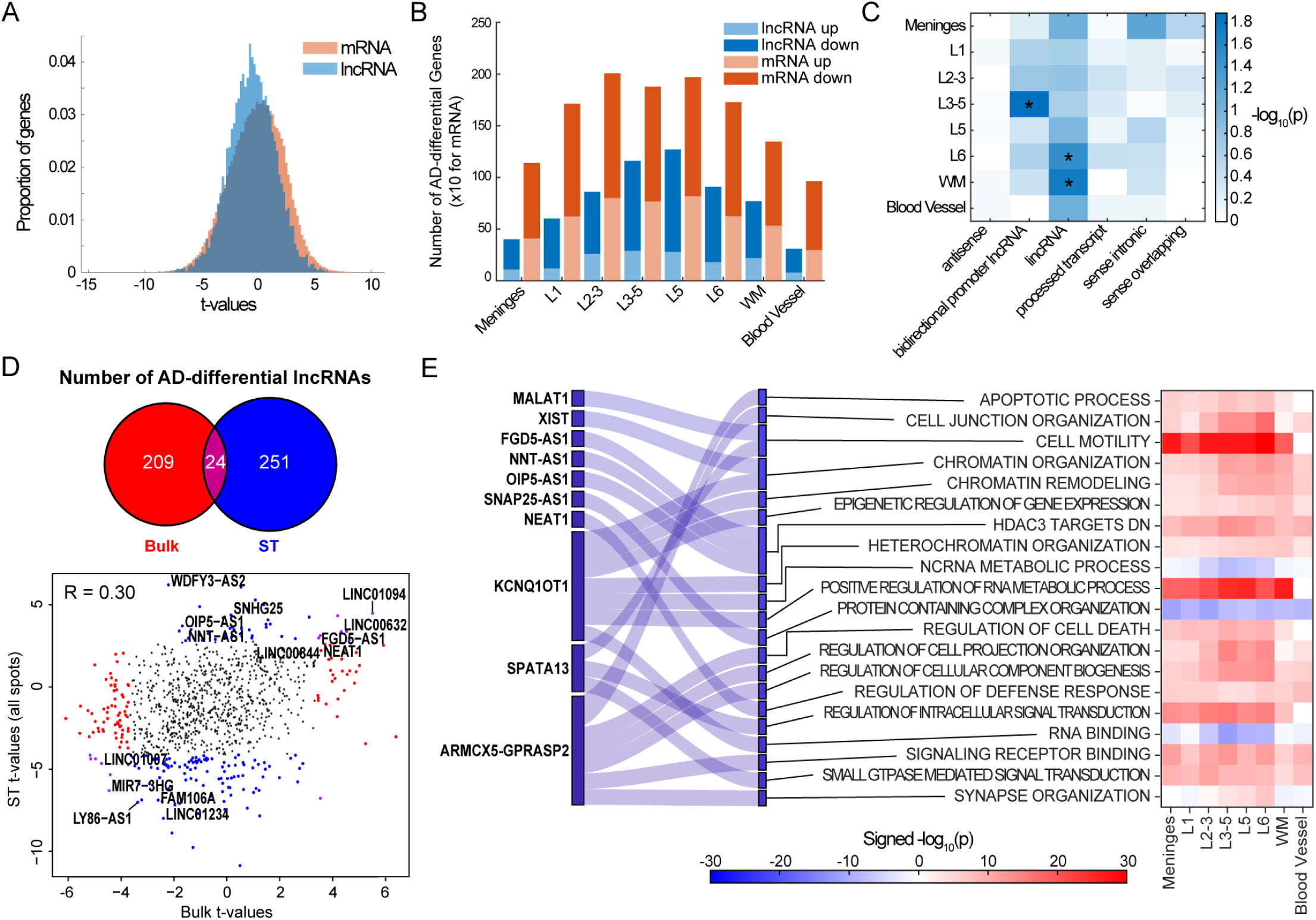
AD-differential lncRNAs and mRNAs identified by ST. **(A)** Histogram showing the distribution of AD-differential t-values for mRNAs and lncRNAs. **(B)** Total numbers of AD-differential lncRNAs and mRNAs for each subregion. **(C)** Heatmap showing the enrichment of AD-differential lncRNAs for specific biotypes (*p<0.05). **(D)** Overlaps in AD-differential lncRNAs and mRNAs between ST and bulk RNA-seq. (bottom) Scatterplot of ST (y-axis) vs. bulk RNA-seq (x-axis) of AD-differential t-values for all co-detected lncRNAs. Correlation value (R) is shown at the top left of the plot. Red dots are genes significant only based on bulk data, blue dots are genes significant based on ST data, and purple dots are genes significant for both data types. **(E)** Sankey diagram and heatmap showing a subset of enriched gene sets and AD differential lncRNAs belonging to those gene sets.

To determine the potential functional roles of these AD DE lncRNAs, we applied gene set enrichment analysis (GSEA) to the set of t-values across all genes for each subregion. Focusing on gene sets comprising AD DE lncRNAs, we observed enrichment for processes, such as cell motility (*MALAT1*, *SPATA13*), apoptosis (*ARMCX5-GPRASP2*), chromatin organization (*XIST*, *KCNQ1OT1*), and regulation of defense responses (*NEAT1*) (Fig. 3E; Table S6-7). Notably, we observed enrichment for HDAC targets, which included three AD DE lncRNAs, namely *FGD5-AS1*, *NNT-AS1*, and *OIP5-AS1* (Fig. 3E; Table S7).

### AD DE lncRNAs revealed by snRNA-seq

We further examined AD DE lncRNAs using the snRNA-seq data (15) and observed that more DE lncRNAs are negatively associated with AD than DE mRNAs (Fig. 4A-B; Table S5), consistent with our ST findings. Additionally, AD DE lncRNAs were enriched for lincRNAs, particularly in inhibitory neurons and oligodendrocyte precursor cells (OPCs) (Fig. 4C). Compared to DE mRNAs, DE lncRNAs displayed greater cell-type specificity in oligodendrocytes, inhibitory neurons, and excitatory neurons (Fig. 4D). Overlaps between subregion-specific and cell-type-specific AD DE lncRNAs were minimal (Fig. 4E). Pairwise t-value correlations revealed moderate alignment between L3–5 and inhibitory neurons as well as between blood vessels and microglia/OPCs (Fig. 4F). While a larger proportion of DE mRNAs were shared across subregions and cell types (Fig. 4E), t-value correlations between subregions and cell types remained weak (Fig. 4G), likely reflecting technological differences between ST and snRNA-seq.

**Figure 4.**
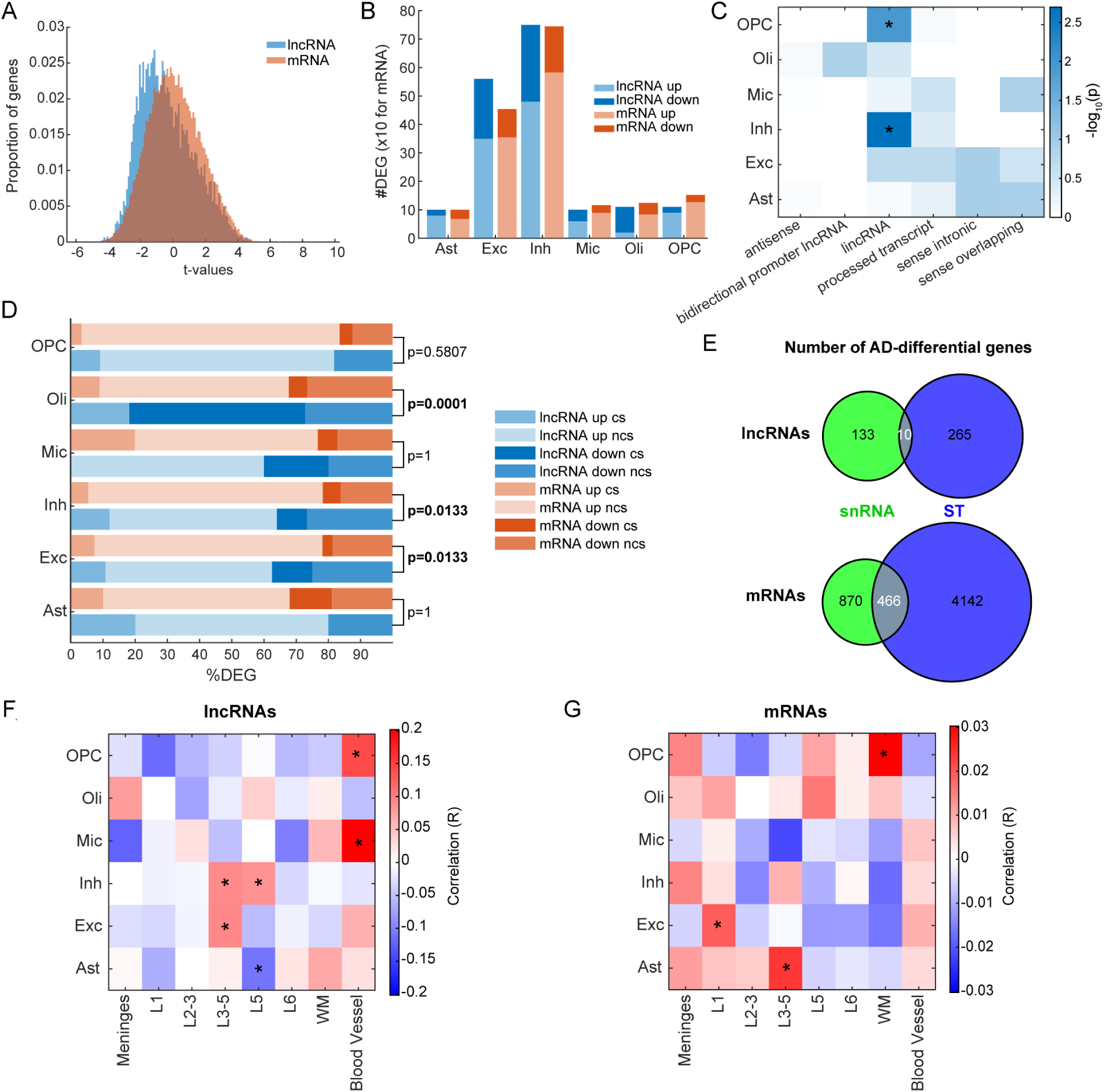
Cross-modality comparison of AD-differential lncRNAs. **(A)** Histogram showing the distribution of AD-differential t-values for mRNAs and lncRNAs based on snRNA-seq data. **(B)** Total numbers of AD-differential lncRNAs and mRNAs for each cell type. **(C)** Heatmap showing the enrichment of cell-type-specific AD-differential lncRNAs for specific biotypes. **(D)** The percentage of DEGs in each cell type, split by lncRNA/mRNA, up/down in AD, and whether they are cell-type-specific (cs) or not (ncs). Fisher’s exact test was applied to assess differences in cell-type-specificity between lncRNA and mRNA with p-values indicated. **(E)** Overlaps in AD-differential lncRNAs and mRNAs between ST and snRNA-seq. **(F-G)** Heatmaps showing correlation (R) between subregions and cell types based on their AD differential t-values (*p<0.05) for lncRNAs **(F)** or mRNAs **(G)**.

### Subregion-resolved AD DE lncRNAs and their cell-type expression pattern

We then examined the subregion specificity of AD DE lncRNAs in our ST data and found DE lncRNAs to be generally more subregion-specific than DE mRNAs, with 115 of 275 DE lncRNAs showing subregion-specific differential expression in the cortex (Fig. 5A; Table S5). These subregion-specific DE lncRNAs, regardless of whether they were up-or downregulated, did not appear to be cell-type-specific (Fig. 5B). The remaining 160 of 275 AD DE lncRNAs were differentially expressed across multiple subregions, with the majority (∼93%) not previously reported (Fig. 5C).

**Figure 5.**
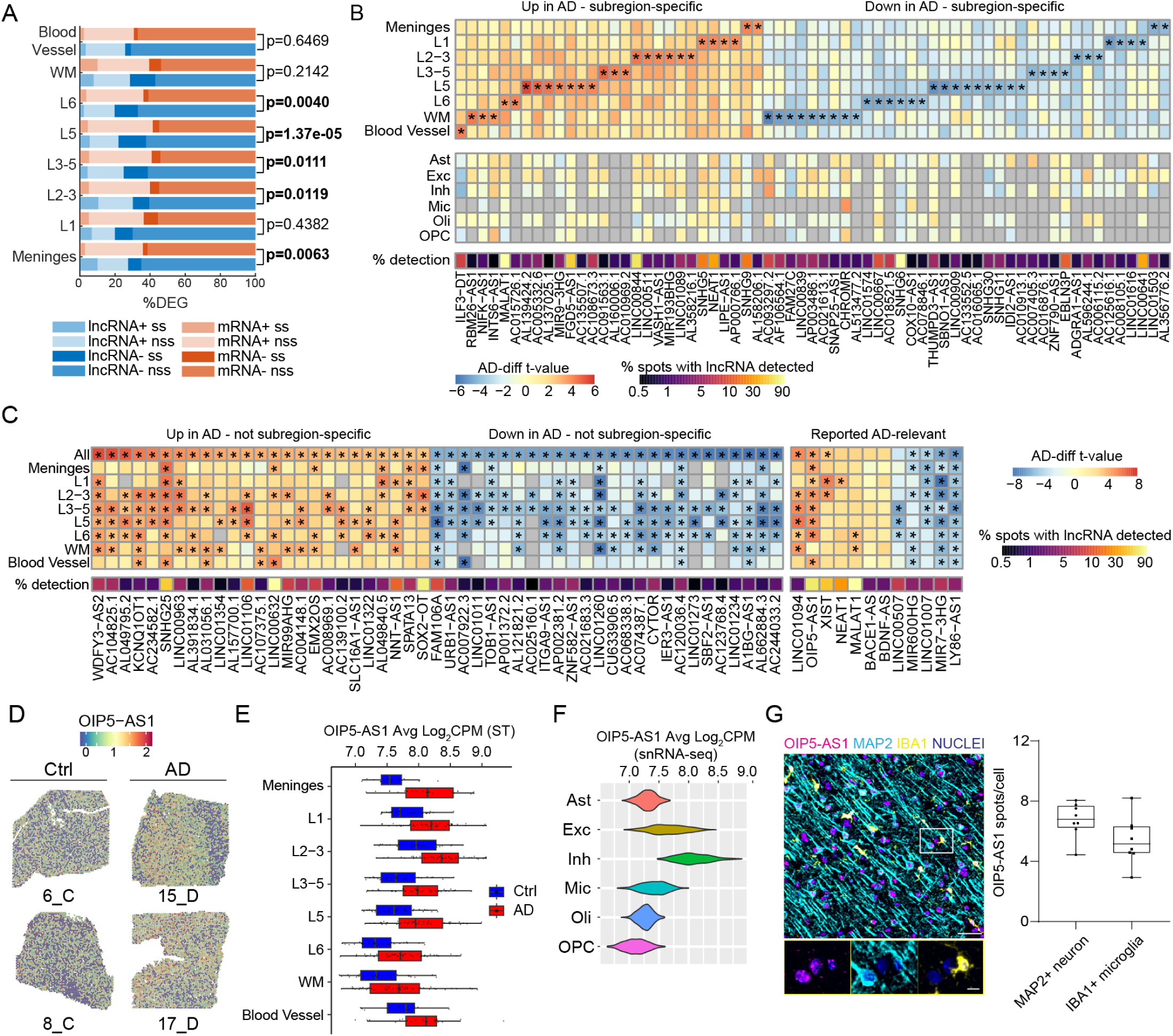
Subregion specificity of AD-differential lncRNAs. **(A)** The percentage of DEGs in each ST subregion, split by lncRNA/mRNA, up/down in AD, and whether they are subregion-specific (ss) or not (nss). Fisher’s exact test was applied to assess differences in subregion-specificity between lncRNA and mRNA with p-values indicated. **(B)** (top) Heatmap showing t-values of AD-differential lncRNAs passing FDR correction specifically for one ST subregion, i.e. subregion-specific (*p<5E-4 in designated subregion and p>0.001 in all others). (bottom) AD-differential t-values for 6 brain cell types. An additional row below shows the percentage of ST spots with non-zero expression for each lncRNA. **(C)** Heatmap showing t-values of AD-differential lncRNAs passing FDR correction in two or more ST subregions, i.e. not subregion-specific (*p<5e-4). An additional row below shows the percentage of ST spots with non-zero expression for each lncRNA. **(D)** *OIP5-AS1* spatial expression plots for two control (Ctrl) and two AD sections. **(E)** Boxplot of subregion-specific pseudobulk expression of *OIP5-AS1* split by AD status. **(F)** Violin plot of cell-type-specific expression of *OIP5-AS1*. **(G)** Combined IHC/RNAScope of human postmortem DLPFC shows that *OIP5-AS1* is expressed in neurons and microglia. Scale bars = 50μm upper, 10μm lower. Boxplot indicates the average number of RNAscope *OIP5-AS1* spots per cell (n=8 individuals).

Some AD DE lncRNAs displayed high expression in one subregion but showed predominant AD effects in other subregions. For example, *LINC00844* was highly expressed in WM/oligodendrocytes (Fig. 2B-C) but exhibited its strongest AD effects in L2-3 (Fig. 5B). Similarly, *LINC00507* showed high expression in L2-3 but demonstrated predominant AD effects in L3-6 (Fig. 5C).

Among previously reported AD DE lncRNAs (9–12,19) (Figure 5C), we observed higher expression of *MALAT1*, *NEAT1, and XIST* in subregion(s) of AD individuals, Interestingly, *MALAT1 has been* reported to suppresses neuronal apoptosis and inflammation and promote neurite outgrowth in AD cellular models(20,21). *NEAT1* induces Aβ-induced neuronal damage and regulates tau phosphorylation and (22,23). Increased *XIST* levels have been previously reported in the brains of female AD patients and linked to X Chromosome Interaction, a process contributing to female-specific vulnerability in AD(24,25) Notably, we observed consistent downregulation of *LY86-AS1* and *MIR7-3HG,* along with upregulation of *OIP5-AS1* in AD across all cortical layers in our ST data (Fig. 5C-E). The functional roles of *LY86-AS1* and *MIR7-3HG* are emerging, with a growing number of studies implicating these lncRNAs in diverse disease contexts. *LY86-AS1* has been linked to type 2 diabetes (26), autoimmune diseases (27), and the progression of multiple myeloma(28), and in AD it shows a negative correlation with Braak stage (*19*)*; MIR7-3HG* has been associated with autophagy regulation and cell proliferation in cancer (29), and in AD it has been related to ferroptosis and immune cell infiltration in postmortem brains(30). By contrast, *OIP5-AS1* has been implicated in a broad range of diseases (31), especially cancer (32–36), where it regulates key cellular processes such as proliferation, cell cycle progression, and apoptosis, and modulation of inflammatory responses. In the nervous system, ischemic injury models show that *OIP5-AS1* modulates neuronal apoptosis and microglia/macrophage inflammatory programs, and is enriched in M2 microglia–derived exosomes (37–39). Recent integrated human data have also identified the *OIP5-AS1/miR-129-5p/CREBBP* axis as a potential therapeutic target for metal toxicity– induced AD pathogenesis (40).

Assessing cell-type expression of *OIP5-AS1* with the snRNA-seq data, we detected *OIP5-AS1* across all major cell types with relatively higher expression in microglia and neurons (Fig. 5F). Orthogonal validation by RNAscope combined with IHC confirmed *OIP5-AS1* signals in IBA1^+^ microglia and MAP2^+^ neurons of ROSMAP DLPFC tissue sections (Fig. 5G).

### *OIP5-AS1* knockdown in iTF-microglia alters gene expression consistent with inflammatory activation

Motivated by above findings, we conducted functional interrogation of *OIP5-AS1* in human iPSC-derived microglia. Using a line with inducible expression of six transcription factors, we generated induced-transcription factor microglia-like cells (iTF-Microglia) via an eight-day protocol (Fig. 6A) (41). On day 8, we transfected cells with two locked nucleic acid (LNA)-modified antisense oligonucleotides (ASOs) targeting *OIP5-AS1* or a non-targeting control and then harvested cells 24 hours later for RNA-seq. *OIP5-AS1* knockdown produced no detectable changes in cell viability or morphology (Fig. 6B). The iTF-microglia expressed the canonical microglial marker IBA1, and the percentage of IBA1-positive cells was similar between unstimulated and ASO-treated conditions (Fig. 6C), indicating that ASO treatment did not impair general microglia identity.

**Figure 6.**
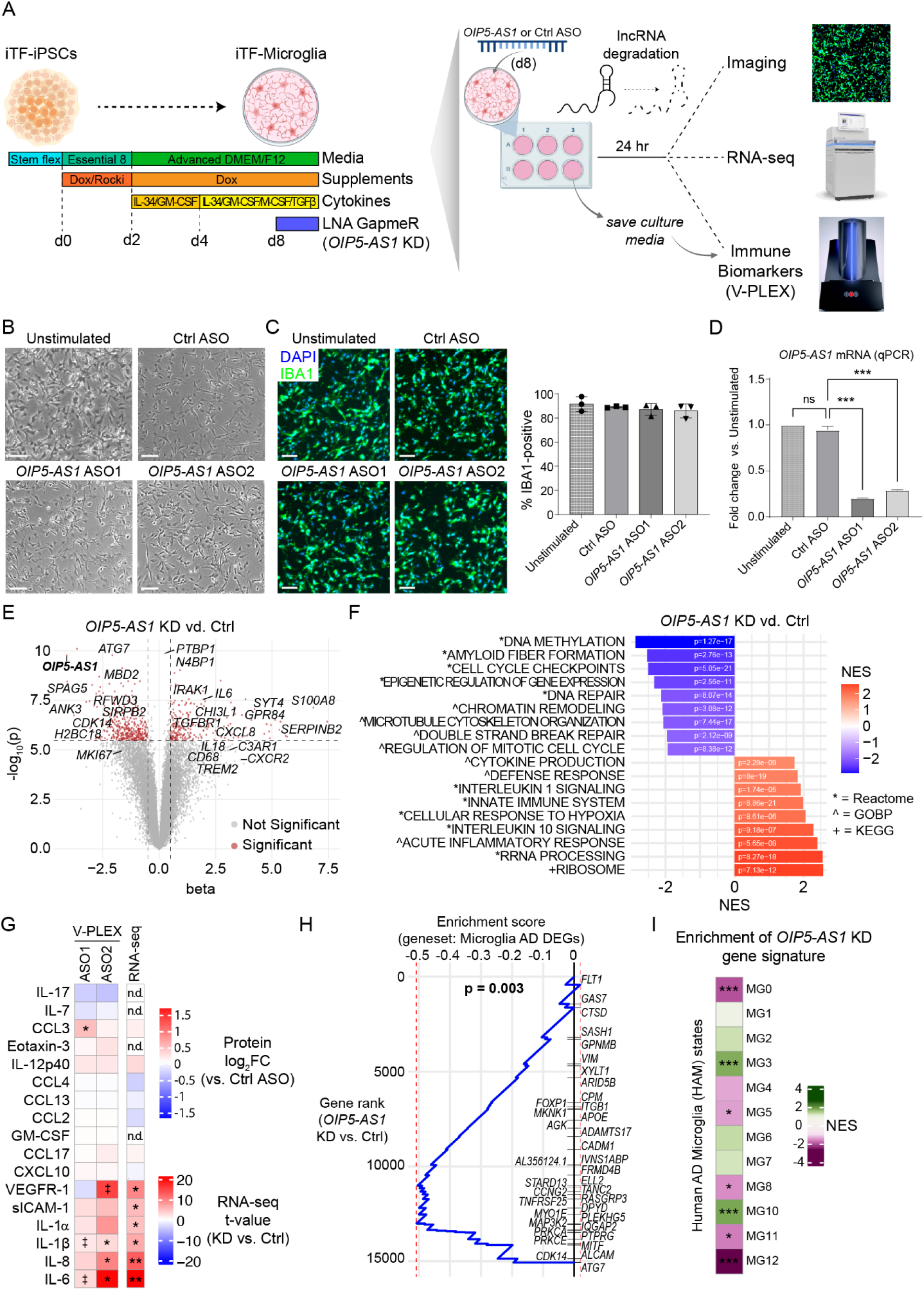
Knockdown of *OIP5-AS1* in cultured microglia uncovers putative roles in AD-relevant biological processes. **(A)** Flowchart of experimental design. Inducible iPSCs were converted to microglia-like cells (iTF-Microglia) using a published protocol (ref), then transfected with ASOs targeting *OIP5-AS1* for degradation (or a non-targeting control ASO). **(B)** Phase contrast images of iTF-Microglia 24 hours post-transfection with ASOs (scale bars = 100 µm). **(C)** Immunofluorescence images of iTF-Microglia 24 hours post-transfection with ASOs (scale bars = 100 µm). Quantification is shown on the right, indicating no significant difference in the percentage of IBA1-positive cells (n=3 biological replicates). **(D)** Quantification of *OIP5-AS1* expression assessed by qPCR (***p<0.001; ns: not significant). **(E)** Volcano plot of RNA-seq results, showing beta and-log10(p) for all genes. Points labeled red have an abs|beta| of > 0.5 and a p<2.2e-06 (Bonferroni FDR). A subset of relevant genes are labelled. **(F)** Barplot showing NES values from GSEA of *OIP5-AS1*-KD RNA-seq for a subset of terms. **(G)** V-PLEX immunoassay results for all analytes detected above the limit of detection, for each *OIP5-AS1* ASO (*p<0.05, キp<0.2). On the right is the RNA-seq t-value (KD vs. Ctrl) for the corresponding gene (n.d.: not detected; *p<2e-4, **p<2e-6). **(H)** functional GSEA of *OIP5-AS1* KD signature compared with microglia-specific AD-differential genes. AD-differential genes that reach dependent FDR (p<3.26e-05) are labelled. **(I)** GSEA NES heatmap showing enrichment of KD signature for human AD microglia (HAM) states.

Reassuringly, *OIP5-AS1* was effectively downregulated following knockdown with either ASO1 or ASO2 (Fig. 6D-E; RNA-seq beta =-4.09, p = 1.69e-10). Transcriptomic profiling of *OIP5-AS1* knockdown (*OIP5-AS1*-KD) cells revealed marked gene expression changes, with upregulation of pathways related to innate immune signaling (*NLRP3*, *TREM1*, *CXCR2*), inflammation (*IL6*, *S100A8*, *ADAM8*), cytokine response (*IRAK1*, *CHI3L1*, *TRIM27*), and rRNA processing (*NOP10*, *MRM1*, *RBM28*) (Fig. 6F; Table S8). Conversely, gene sets involved in mitotic control (*BIRC5*, *PLK1*, *KIF2C*), DNA methylation (*DNMT1*, *H4C14*, *H3C4*, and other histone genes), and DNA repair (*MLH1*, *BRIP1*, *RPA1*) were downregulated, accompanied by a strong repression of canonical histone genes (50 of the 69) (Fig. S2A). These findings align with previously reported functional roles of *OIP5-AS1* in cell cycle regulation and inflammation in cancer and other diseases (31,42,43). Additionally, *OIP5-AS1* knockdown also resulted in downregulation of multiple genes involved in amyloid fiber formation, including *APP*, *ADAM10*, and *SORL1* (Fig. 6F; Table S8), indicating a potential role of *OIP5-AS1* in amyloidogenesis.

### *OIP5-AS1* knockdown in iTF-microglia alters cytokine secretion

To evaluate the impact of *OIP5-AS*1 knockdown on immune signaling at the protein level, we performed a Meso Scale multiplex immunoassay (‘V-PLEX”) on culture supernatants to quantify soluble cytokines and chemokines 24 hours after transfection. Although variability between biological replicates was high and many analytes were near the limit of detection, a few showed reproducible changes across experiments. In particular, IL-6, IL-1β, IL-8 (*CXCL8*) and CCL3, all pro-inflammatory cytokines/chemokines, were elevated in *OIP5-AS1*-KD cells compared to controls (Fig. 6G). Several additional factors—including sICAM-1, VEGFR-1 (*FLT1*), and IL-1α—showed trends toward increased secretion, though differences did not reach statistical significance (Fig. 6G). To assess potential impairments in phagocytosis, we incubated control and *OIP5-AS1*-KD cells with pHrodo-labelled amyloid-beta (Aβ1-42) for two hours, then assessed Aβ uptake by flow cytometry. No differences were observed, with all groups phagocytosing ∼90-96% of Aβ (Fig. S2B).

### *OIP5-AS1*-KD genes are associated with AD microglia gene signatures

To evaluate whether the *OIP5-AS1*-KD transcriptional signature reflects AD-related microglia programs, we performed GSEA using a set of microglia-specific AD DE genes (15). This analysis revealed a negative enrichment (p=0.003), indicating that microglia AD genes were on average downregulated upon *OIP5-AS1* knockdown (Fig. 6H). We next compared the *OIP5-AS1-*KD signature to reported human AD microglia states (44) (Fig. 6I). We found positive enrichment for MG3 (ribosome biogenesis; disease-associated microglia) and MG10 (a pro-inflammatory state), and negative enrichment for MG0 (homeostatic) and MG12 (cycling) (Fig. 6I). These findings suggest that *OIP5-AS1* knockdown shifts microglia gene programs toward inflammatory and disease-associated states while reducing homeostatic and cycling programs.

## Discussion

Recent advancements in sequencing technologies have revealed that a large portion of the mammalian genome is transcribed into lncRNAs (1). Leveraging published ST (14) and snRNA-seq datasets (15), we mapped the spatial and cellular expression of lncRNAs in DLPFC of aged human brains. We found that lncRNAs exhibit greater subregion-specific expression than mRNAs and subregion-enriched lncRNAs are generally not cell-type specific, and vice versa. These observations likely reflect the heterogeneity of cell types within each subregion and the shared developmental origins of cell types that span cortical layers. Nonetheless, the presence of subregion-specific and cell-type-specific lncRNAs suggests that some lncRNAs function within precise spatial regions to support localized processes, while others serve the specific needs of individual cell types.

For AD DE lncRNAs, we observed a moderate correlation between those identified from ST and bulk RNAseq data (Fig. 3D), but only minimal overlap with results from snRNAseq (Fig. 4E). This discrepancy may reflect differences in sequencing platforms as well as general technical limitations in detecting lncRNAs. Whether subregion-specific or not, AD DE lncRNAs in ST data are often expressed across multiple major cell types, suggesting that they may participate in multicellular and diverse processes. Consistently, the lncRNA-enriched modules are enriched for diverse biological processes (Fig. S1). In addition, gene set enrichment analyses (Fig. 3E) indicate the potential involvement of AD DE lncRNAs in a wide range of regulatory processes, including epigenetic regulation, chromatin remodeling, RNA metabolism, apoptosis, cell motility, regulation of cell death, and synaptic organization.

Among previously reported AD DE lncRNAs (9–12,19), *OIP5-AS1* is consistently upregulated across cortical subregions in our ST data from AD brains (Fig. 5C-E). Prior animal studies in brain ischemic injury have shown that *OIP5-AS1* plays a pivotal role in modulating neuronal apoptosis, pyroptosis, and microglia/macrophage activation and is enriched in M2 microglia–derived exosomes (37–39). In our study, antisense oligonucleotide–mediated knockdown of *OIP5-AS1* in iPSC-derived microglia induced robust upregulation of immune and inflammatory pathways (e.g., innate immune signaling, cytokine production, acute inflammatory response) while markedly suppressing programs essential for cell cycle regulation, chromatin maintenance, DNA repair, and histone expression. This dual effect mirrors diverse roles of *OIP5-AS1* in other disease contexts and, importantly, recapitulates differential gene expression patterns observed in maladaptive AD microglial states. Notably, recent integrated human data identify the *OIP5-AS1/miR-129-5p/CREBBP* axis as a potential therapeutic target in metal toxicity– induced AD pathogenesis (40). Together, these findings raise the possibility that modulating *OIP5-AS1* activity could shift microglial states toward neuroprotection, providing a potential avenue for therapeutic intervention in AD and related neurodegenerative diseases.

## Conclusions

While the brain expresses more tissue-specific lncRNAs than most other tissues (45,46), their spatial and cellular contexts in the human brain remain poorly understood. By leveraging our previously published ST data, along with bulk and snRNA-seq data from ROSMAP participants, we mapped the spatial and cellular expression of lncRNAs in brain tissue and identified spatially resolved and cell-type-specific differentially expressed lncRNAs in AD. Our analysis lays the groundwork for future investigations using more advanced technologies to examine lncRNA expression with greater spatial and cellular resolution. Our functional perturbation experiments on *OIP5-AS1* further support the idea that lncRNAs can play important roles in regulating cellular functions and may contribute to disease processes. Given the relatively limited attention this area has received, our results underscore the need for continued research to fully explore the functional and mechanistic roles of lncRNAs in Alzheimer’s disease and related brain disorders.

## Methods

### Tissue collection for Spatial Transcriptomics

The human brain tissues were obtained through the ROSMAP (Religious Orders Study and Rush Memory and Aging Project) cohort studies at the Rush Alzheimer’s Disease Center, Rush University Medical Center (RUMC), Chicago, IL (47). Both studies were approved by the Institutional Review Board of RUMC. Written informed consent was obtained as was a repository consent and an Anatomic Gift Act, in compliance with national ethical guidelines. We selected 13 female individuals clinically diagnosed as AD and 8 controls with no cognitive impairment (NCI) based on criteria as previously reported (48–50). The brain tissue of those individuals had RIN > 5 and minimal freezing artifacts. Frozen brain blocks of the right-hemisphere DLPFC were cut (∼ 1 cm^3^), transferred on dry ice, and stored at-80°C.

Frozen brain blocks were embedded in OCT and cryosectioned coronally to a thickness of 10 μm using a Leica CM1950 cryostat set at-17°C. Sectioning was done in sets of three (middle section for ST and two adjacent sections for IF), with 10-50 μm between each set. Sections were attached to Visium slides and stored at-80°C in a slide mailer for up to one week before proceeding with spatial transcriptomics. ST-adjacent sections were attached to pre-labeled Leica superfrost plus slides and stored at-80°C in slide mailers for up to two weeks before proceeding with immunohistochemistry.

### Spatial Transcriptomics

Spatial Transcriptomics was performed according to the Visium Spatial Gene Expression User Guide (10X Genomics; CG000239). Briefly, cryosectioned tissue on Visium slides was transferred from - 80°C to the lab on dry ice and quickly thawed for 1 min on a heat block at 37°C, then immediately transferred to methanol pre-equilibrated to-20°C and fixed at-20°C for 30 min. Sections were then stained with hematoxylin and eosin, and brightfield images were acquired with a Nikon Ti2 with NIS-Elements AR 5.11.01 64-bit software. After imaging, tissues were immediately permeabilized in 0.1% pepsin diluted in 0.1 M HCl for 18 min at 37°C. Reverse transcription, second-strand synthesis, cDNA amplification (15 cycles), and library construction were performed according to the manufacturer’s instructions. For the first four individuals (#2, 8, 12, and 18), protocol version’Rev A’ was followed, and the sample index PCR was 12 cycles. For the remaining samples, protocol version’Rev D’ was followed, and the sample index PCR was 13-14 cycles. Library concentration was quantified by Qubit 1x dsDNA HS using a SpectraMax M3 plate reader. Library concentration and size distribution were assessed by Fragment Analyzer (Agilent). Sequencing was performed in three batches on an Illumina NovaSeq, targeting a minimum of 50,000 raw reads per spot.

### Spatial transcriptomics sequencing data processing

Sequencing data were pre-processed with the Space Ranger pipeline (10x Genomics), which mapped the barcodes against the human genome (Gencode 32), generated FASTQ files, and generated a count matrix for each section. The individual files were then merged into a Seurat object (Seurat v4.2.0) and were processed in R. We calculated per-spot quality metrics and spots with low-measured genes (< 500) were removed. The Seurat::SCTransform function was used to normalize and scale the UMI counts based on regularized negative binomial regression. Principal component analysis (PCA) was performed on the top 3000 variable genes. Using the top 50 PCs and accounting for technical and biological covariates such as age, RIN, library batch, and donor, we integrated the datasets using the Harmony package (v0.1.0). We explored various tools for data integration and chose Harmony for its high performance, usability, and scalability. We only used Harmony integration to identify clusters corresponding to brain regions. All downstream analyses (differential gene expression, cell proportion estimates, and ligand-receptor analysis) were performed on pre-integration count data. After visualizing the standard deviation attributed to each Harmony embedding as an elbow plot, we decided to use the top 10 Harmony embeddings. We applied Seurat::runUMAP to compute UMAP (uniform manifold approximation and projection) dimension reduction and Seurat::findNeighbors and Seurat::findClusters to identify clusters based on a shared nearest neighbor (SNN) clustering algorithm (Louvain clustering with resolution set to 0.3; all other parameters default). This resulted in 9 clusters, which were annotated based on their canonical markers and spatialLIBD (v7.5) to the closest anatomical cortical layer (L1, L2-3, L3-5, L5, L6), white matter (WM), meninges, and specific cellular subpopulations, namely cells with high expression of hemoglobin genes (’Blood Vessel’) and SST/NPY (’Interneuron’). The Interneuron cluster had too few spots for reliable pseudobulking so was not analyzed further.Seurat::FindMarkers was used to identify cluster-enriched (e.g. subregion-specific) lncRNAs reported in Fig. 2C and Table S1. For Fig. 2C, a log2FC cutoff of 0.1 was employed, resulting in 50 lncRNAs.

### Co-expression network analysis in spatial transcriptomics

The modules of co-expressed genes were derived using the Speakeasy (SE) algorithm v1, Matlab version (17). We created pseudo-bulk gene expression by summing the counts per donor, then splitting by ST spot cluster. Lowly expressed genes (less than 1 count per million, CPM) were removed, and the variancePartition R function was applied to capture the major sources of variation, as previously described (14). The input for SE consisted of the residuals after normalization using tmm.voom and adjustment for library batch, RIN, age, and post-mortem interval (PMI). We built the modules separately for each ST spot cluster, using the results from 100 initializations of SE and Spearman correlation. This resulted in 193 modules, each with at least 30 genes (L1 = 30, L2-3 = 21, L3-5 = 20, L5 = 20, L6 = 20, Meninges = 33, WM = 20, and Blood Vessels = 29). The module assignments are provided in Table S2.

### Functional Enrichment Analysis (FEA) of Modules

Each module was annotated using the gprofiler software (v2_0.2.1) via the g:GOSt R function (51). This tool maps genes to functional information sources and detects statistically significantly enriched terms. These sources include the GO database, pathways from KEGG, Reactome, and WikiPathways, miRNA targets from miRTarBase, regulatory motif matches from TRANSFAC, tissue specificity from the Human Protein Atlas, protein complexes from CORUM, and human disease phenotypes from the Human Phenotype Ontology. For multiple testing correction, we used the default g:SCS method unless otherwise specified. The FEA results are available in Table S4.

### Module Preservation (MP) Analysis

To assess whether the modules were preserved across distinct ST spot clusters, we used the MP function from the WGCNA R package (v1.70-3) (52). Pairwise comparisons were performed for all 193 modules. We then estimated the mean and variance of the preservation statistic under the null hypothesis that there is no relationship between module assignments in the reference and test datasets. Fifteen distinct preservation statistics were calculated (e.g., module density, mean expression, connectivity), each standardized by its mean and variance to define a Z statistic for each preservation measure. Under these assumptions, each Z statistic follows a normal distribution if the module is not preserved. To compare preservation across modules, we calculated a composite preservation measure using the following equation:

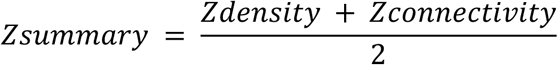

Modules with Z < 2 were considered not preserved, those with Z between 2 and 10 were considered preserved, and modules with Z > 10 were considered highly preserved. Zsummary stats are available in Table S3.

### Statistical analysis to identify AD-differential genes

For each brain section of a subject, we summed the UMI counts of all spots belonging to a subregion to generate a pseudobulk estimate for each gene. We only kept expressed genes with counts per million (CPM) > 1 in at least 80% of the brain sections, resulting in 1,578 lncRNAs and 13,808 mRNAs. With the pseudobulk estimates, we estimated the library sizes of the brain sections using the trimmed mean of M-values (TMM) of edgeR(53) (edgeR_3.36.0) and transformed the pseudobulk estimates into log2 CPM using voom-limma (limma_3.50.1). For each subregion, we excluded brain sections with <50 spots. To contrast the normalized pseudobulk expression between AD and controls, we applied linear mixed models (LMM), accounting for age, PMI, RIN, and batch as fixed effects and modeled within-subject correlation with a random effect. We declared significance at an *α* of 0.05 with false discovery rate (FDR) correction across all genes and all subregions.

For interpretation, we applied GSEA (54) with AD-differential t-values of each subregion as scores, and GO terms and canonical pathways as gene sets. To characterize lncRNAs with respect to brain cell types, we used the pseudobulk expression data from a published snRNAseq dataset of 424 ROSMAP individuals (15). For each of the major cell types, we contrasted pseudobulk expression of AD subjects vs. controls using multiple regression, accounting for age, PMI, RIN, and batch. For further comparison, we repeated this AD-differential expression analysis with bulk DLPFC RNA-seq data from 1049 individuals (16).

### RNAscope and immunofluorescence co-staining of postmortem tissue

Human FFPE DLPFC sections were stained with neuron (MAP2; Sigma #M2320-100UL) and microglia (IBA1; Wako #019-19741) markers followed by RNAscope with Hs-OIP5-AS1-O1-C1 probe (ACD #1559738-C1) using a Bond-Rx research stainer (Leica Biosystems) using an Opal 6-plex detection kit (Akoya Biosciences #NEL871001KT). Briefly, sections were baked for 30 min at 60°C prior to dewaxing for 30 s at 72°C. Sections were antigen retrieved using HIER with a pH 6.0 citrate buffer solution (Leica Biosystems #AR9961) for 20 min at 100°C followed by 10 min peroxide block. MAP2 (1:100) was incubated for 30 min followed by 10 min Opal polymer HRP Ms+Rb and 10 min Opal signal development with Opal 520 diluted at 1:100. Sections were antigen retrieved for 20 min in ph 6.0 buffer again and the process is repeated with IBA1 (1:500) paired with Opal 690 (1:150). The sections were then stained with RNAscope probe against *OIP5-AS1* using RNAScope LS Multiplex Fluorescent Reagent Kit (ACD #322800) according to the manufacturer with the following modifications. The sections were antigen retrieved using HIER with a pH 6.0 buffer for 20 min at 100°C, washed, and treated with ACD Protease III for 15 min at 40°C prior to probe hybridization for 2 h at 40°C. Opal 620 (1:500) was used for signal development. Slides were counter stained with 10X spectral DAPI for 5 min before mounting. Imaging was performed using Phenocycler-Fusion 2.0 (Akoya Biosciences) at 40X. Images were spectrally umixed using inForm (Akoya Biosciences) and composite TIFs exported to QuPath (v0.3.2) for quantitative analysis. Cells were defined in each ROI using QuPath Cell Detection tool with the following parameters, requested pixel size = 0.5μm, background radius = 8μm, median filter radius =0μm, Sigma value for Gaussian filter = 1.5μm, minimum nucleus area = 10μm2, maximum nucleus area =400μm2, and intensity threshold = 10. Cell parameters include a cell expansion of 5 μm from the cell nucleus and general parameters include smoothing of the detected nucleus/cell boundaries. To quantify OIP5-AS1, subcellular spot parameters were set to detection threshold = 0.08, split by intensity and shape to true, expected spot size = 0.05μm2, min spot size = 0.01μm2, max spot size =1μm2 and include clusters set to true. For cell type specific expression, cells were thresholded on their MAP2 and IBA1 mean cell intensity by section and cells of ambiguous identity were not included in the analysis.

### Human iPSC-derived microglia and *OIP5AS1* knockdown

The iTF-iPSC line (SF2021-173 from Kampmann lab’s resource website) is a generous gift from the Kampmann lab (41). Microglial differentiation was performed in 6-well or 24-well plates as described above with the exception that the day 8 media change was performed on day 7 of differentiation. For *OIP5-AS1* knockdown, antisense LNA GapmeRs were designed using GeneGlobe and tested in cultured cells for knockdown compared to negative control A oligo (Qiagen). GapmeRs were resuspended in TE at 0.5 µg/µl and stored at-20°C. On day 8 of differentiation, GapmeRs were thawed and combined with FuGENE HD (Promega) in Opti-MEM at a ratio of 1µl FuGENE:1µl GapmeR:25µl Opti-MEM. Solutions were vortexed briefly and allowed to settle for 10 min before transfection. Microglia in each well were treated with 2µg GapmeR for 24 h. On day 9, cells were washed briefly in PBS, and RNA extraction (RNeasy, Qiagen) was performed. Samples from three biological replicates were used for RNA-seq or cDNA synthesis (High-Capacity cDNA Reverse Transcription Kit, ABI). Quantitative PCR assays were performed with the Taqman probe (Hs03677189_g1, ThermoFisher) normalized to GAPDH.

### Mesoscale Multiplex Immune Biomarker Assay

Conditioned media from day 9 culture was collected and frozen at-20°C. To simultaneously assess multiple immune biomarkers, conditioned media was thawed on ice, processed, and analyzed using the V-PLEX Human Biomarker 39-Plex Kit (Mesoscale, K15209D-1) per the manufacturer’s protocol. Raw data was processed using MSD Discovery Workbench (v4) software and statistical analysis was performed using GraphPad Prism 10. Comparisons between groups utilized one-way ANOVA followed by Tukey’s *post hoc* test and corrected p-values for multiple comparisons were reported.

### Amyloid beta Labeling and Phagocytosis Assay

Amyloid-beta (Aβ1-42) was reconstituted and aggregated according to the Fujufilm/Cellular Dynamics Labeling Amyloid Beta with pHrodo Red protocol. All reagents required to reconstitute Aβ were provided in the aggregation kit (rPeptide; Cat. A-1170-2). Briefly, lyophilized Aβ was reconstituted in 250 µL 10 mM NaOH and transferred to a 1.5 mL microcentrifuge tube. A total of 250 µL HPLC water was added, followed by 56 µL 10x TBS (pH 7.4). The reconstituted Aβ was allowed to aggregate in an incubator overnight under normoxic conditions. After overnight incubation, the aggregates were collected by centrifugation at 16,000 x g for 1 min at room temperature. Supernatant was removed and aggregates were washed with 200 μL fresh 0.1 M NaHCO3 in water. A total of 1 mg pHrodo Red, Succinimidyl Ester (ThermoFisher) was reconstituted in DMSO to a final concentration of 10 mM immediately before use according to the manufacturer’s protocol. pHrodo Red dye was added to supernatant-free AB aggregates and mixed well. The reaction was incubated in the dark at room temperature for 1 hour. The reaction tube was centrifuged as described and supernatant removed. To remove excess dye, 1 mL of cold methanol was added to the aggregates, and the sample was vortexed on high speed for 10 seconds. The reaction tube was centrifuged again and the supernatant removed. Aggregates were washed with 1 mL of HBSS, vortexed on high for 10 seconds, and centrifuged before removing the supernatant. Aggregates were washed an additional 3 times with HBSS as described. Finally, the aggregates were resuspended in 200 uL HBSS, sonicated for 10 min, and stored at-20 °C in individual aliquots.

On day 8 of differentiation, cells in 6-well plates were treated with GapmeRs or controls as described above. On day 9, 24 hrs after treatment, Aβ was added at a dilution of 1:200 to each well for 2 h during which cells were returned to a 37°C incubator. During the last 15 min of Aβ incubation, cells were treated with Calcein Green AM (1:1000, Invitrogen). Media from each well was removed, and cells were washed once with PBS and then treated with 1 mL of TrypLE Express. Plates were returned to the incubator for 7-10 min to detach adherent cells. Cells were collected into their respective round-bottom tube and the wells were washed with 1 mL of warm DMEM/F12 media to collect any remaining cells. Cells were then centrifuged at 300 x g for 5 min at 4°C and supernatant discarded. FACS buffer (2% BSA) was prepared by diluting 30% BSA (Fisher Scientific; Cat. No. 50-203-6474) in DPBS and was used to resuspend the pelleted cells. Resuspended cells were filtered through a 70 µm mesh filter immediately before data was collected by flow cytometry using the Sony SH800 Cell Sorter. Analysis was performed with FlowJo v10. Single cells were gated and selected for analysis based on viability, size, and complexity. The experiment was repeated for a total of three biological replicates.

### Immunostaining of iPSC-derived microglia

iPSCs were differentiated as described above in 24 well plate format and transfected on day 8 with 0.4 µg per well. On day 9, samples were washed in PBS and treated with 4% paraformaldehyde (PFA; Electron Microscopy Sciences, 15714-S) for 15 min. Samples were then washed three times in PBS and stored at 4°C. Cells were permeabilized and blocked in 5% normal donkey serum (Jackson ImmunoResearch, 017-000-121) with 0.02% Triton X-100 (Fisher, BP151-100) in PBS for 1 h at room temperature. Cells were then incubated with a primary antibody (anti-IBA1 1:1000) diluted in a blocking buffer for 2 h at room temperature. Cells were washed three times with PBS and then incubated with secondary antibodies diluted 1:1000 in a blocking buffer for 45 min at room temperature in the dark. Cells were washed three times with PBS and then stained with Hoechst 33342 (1:2000 in PBS; Invitrogen, H1399) for 3 min at room temperature in the dark. Cells were washed one additional time with PBS and imaged using a Nikon Eclipse Ti2-E epi-fluorescent microscope. Quantification of IBA1-positive cells was done in GraphPad Prism.

### RNA-seq of iPSC-derived microglia

RNA was extracted using the RNeasy kit (Qiagen). RNA concentration was determined using Qubit broad range RNA assay (Invitrogen, Q10211) according to the manufacturer’s instructions. 500 ng total RNA was used for RNA-Seq library generation using the Stranded Total RNA Prep with Ribo-Zero Plus (Illumina, 20040529). A Zephyr G3 NGS workstation (Perkin Elmer) was utilized to generate sequencing libraries. Library size and concentrations were determined using an NGS fragment assay (Agilent, DNF-473) and Qubit 1X dsDNA assay (Invitrogen) respectively, according to the manufacturer’s instructions. Libraries were normalized for molarity and sequenced on an Illumina NovaSeq 6000 platform, generating an average of 67.50 ± 7.80 million paired-end reads per sample (2 × 150 bp). Transcript abundance was quantified using kallisto (v0.45.0) with GENCODE v43 gene annotation, producing estimated counts matrices at transcript and gene level. With the gene level count matrix, we first converted it into log2(counts per million (CPM)). For each gene, we fitted a regression model with expression level as outcome and the two KD conditions and two control conditions (coded as four binary variables) as covariates, and contrasted the average effect of KD against control. We then applied GSEA to the resulting set of t-values.

## Declarations

### Ethics approval and consent to participate

The human brain tissues were obtained through the ROSMAP (Religious Orders Study and Memory and Aging Project) cohort studies at the Rush Alzheimer’s Disease Center, Rush University Medical Center (RUMC), Chicago, IL., in compliance with national ethical guidelines. Both studies were approved by the Institutional Review Board of RUMC.

## Consent for publication

Written informed consent was given by the donors for brain autopsy and for the use of material and clinical data for research purposes, as was a repository consent and an Anatomic Gift Act.

## Availability of data and materials

Subregion-and cell-type-enriched lncRNAs, ST gene expression modules, AD-differential genes, and knockdown-dysregulated genes are provided as supplemental tables. Raw and normalized count matrices of the ST data are available at Synapse under accession code syn53141470.

## Author contributions

B.N., D.R.A., K.P.L., S.T., R.A.V., T.C., and A.I. performed data analysis. D.R.A. and S.D.T. conducted ST experiments and sequencing. J.X. performed ST data processing. M.F., V.M., and P.L.D. provided snRNA-seq data. H.V. and N.A.K. conducted RNAScope and IHC experiments. Y.W., B.N., D.R.A., and K.P.L. wrote the manuscript. F.S., D.S., and A.K.L. performed the microglia culture and knockdown experiment. D.A.B. runs the parent ROSMAP study and provides critical input. Y.W. conceived and directed this study, overseeing research activities and leading manuscript preparation. All authors reviewed and approved the final manuscript.

## Acknowledgments

We are grateful to those who agreed to donate their brains for research. We thank all the employees at RADC for their support and assistance. We thank Dr. Martin Kampmann at UCSF for sharing the iTF-iPSC cell line.

## Authors’ Information

Further information and requests may be directed to the corresponding author Yanling Wang

## Materials Availability

This study did not generate new unique reagents.

## Supplementary Figures

**Figure S1.**
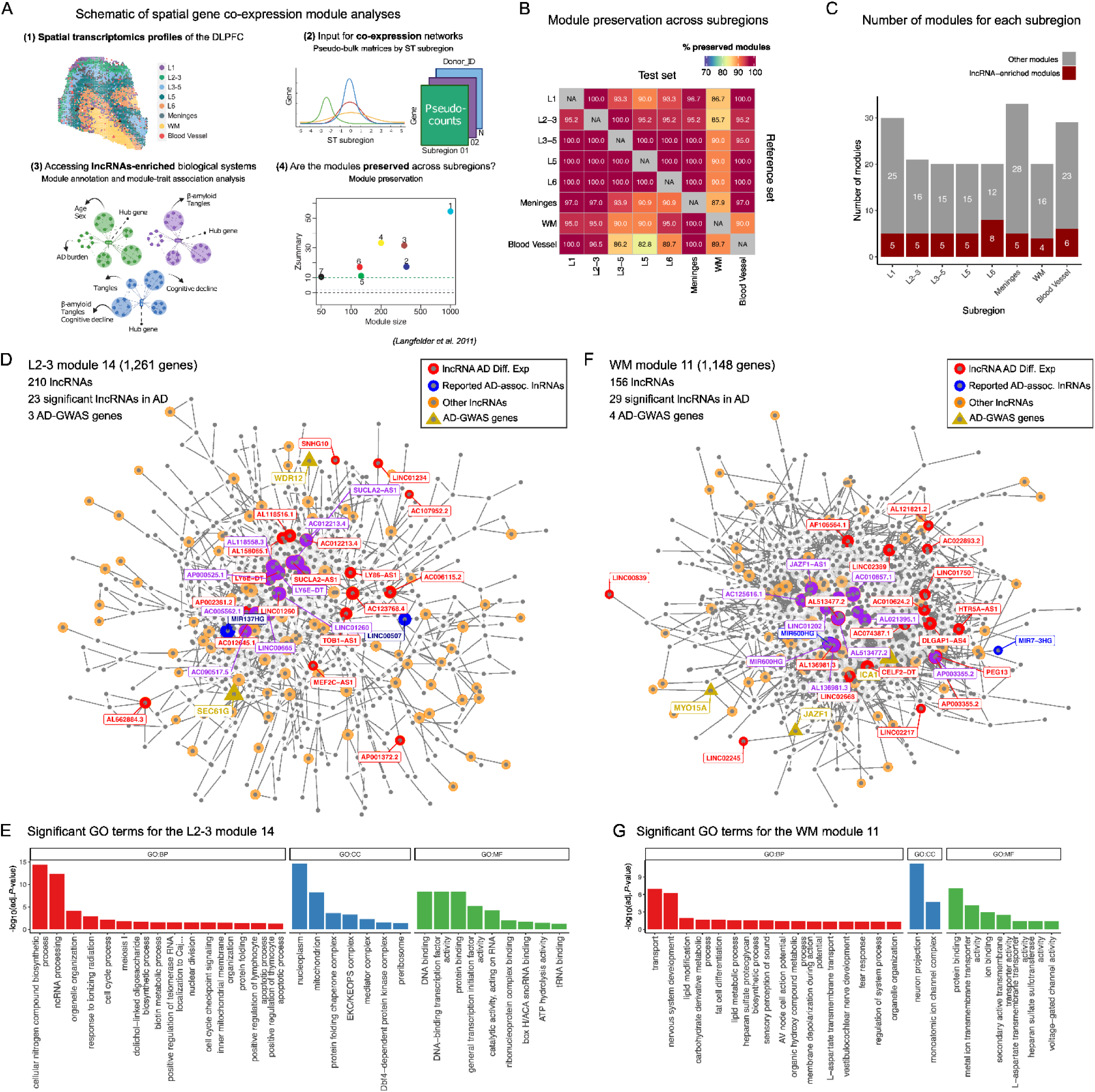
Subregion-specific gene modules. **(A)** Schematic of gene expression module generation and downstream analyses. **(B)** Heatmap showing the percentage of preserved modules between each pair of subregions. Cortical layer modules were more preserved than those derived from WM, blood vessels, and meninges. **(C)** Stacked barplot shows the number of modules in each subregion. Red colors for the lncRNA-enriched modules. Within individual subregions, 15–60% of the modules were lncRNA-enriched. **(D)** Network connectivity map for L2-3 m14, illustrating high connectivity of lncRNAs (correlation > 0.9). This module was enriched with 210 lncRNAs (Bonferroni p < 0.05), including 4 reported AD-differential lncRNAs(9,10,55), 3 AD-GWAS genes(56), and 23 AD-differential lncRNA genes based on our ST analysis (Tables S2, S6). **(E)** Enriched Gene Ontology (GO) terms (FDR at 0.05) from functional enrichment analysis of an example lncRNA-enriched module (L2-3 m14). **(F)** Network connectivity map for WM module 11, illustrating high connectivity of lncRNAs (correlation > 0.95). This module was enriched with 156 lncRNAs (Bonferroni p < 0.05), including 2 previously reported AD-differential lncRNAs(9,10,55), 4 AD-GWAS genes(56), and 29 AD-differential lncRNAs based on our ST analysis (Tables S2, S6). **(G)** Enriched GO terms (FDR at 0.05) from functional enrichment analysis for the WM module 11. **(E, G)** The GO biological process terms were collapsed, keeping only the top GO parent functions to avoid redundancy. The y-axis and bar size represent the significance in-log10(Adj. p-Value). GO biological process (BP), cellular component (CC), and GO molecular function (MF).

**Fig. S2.**
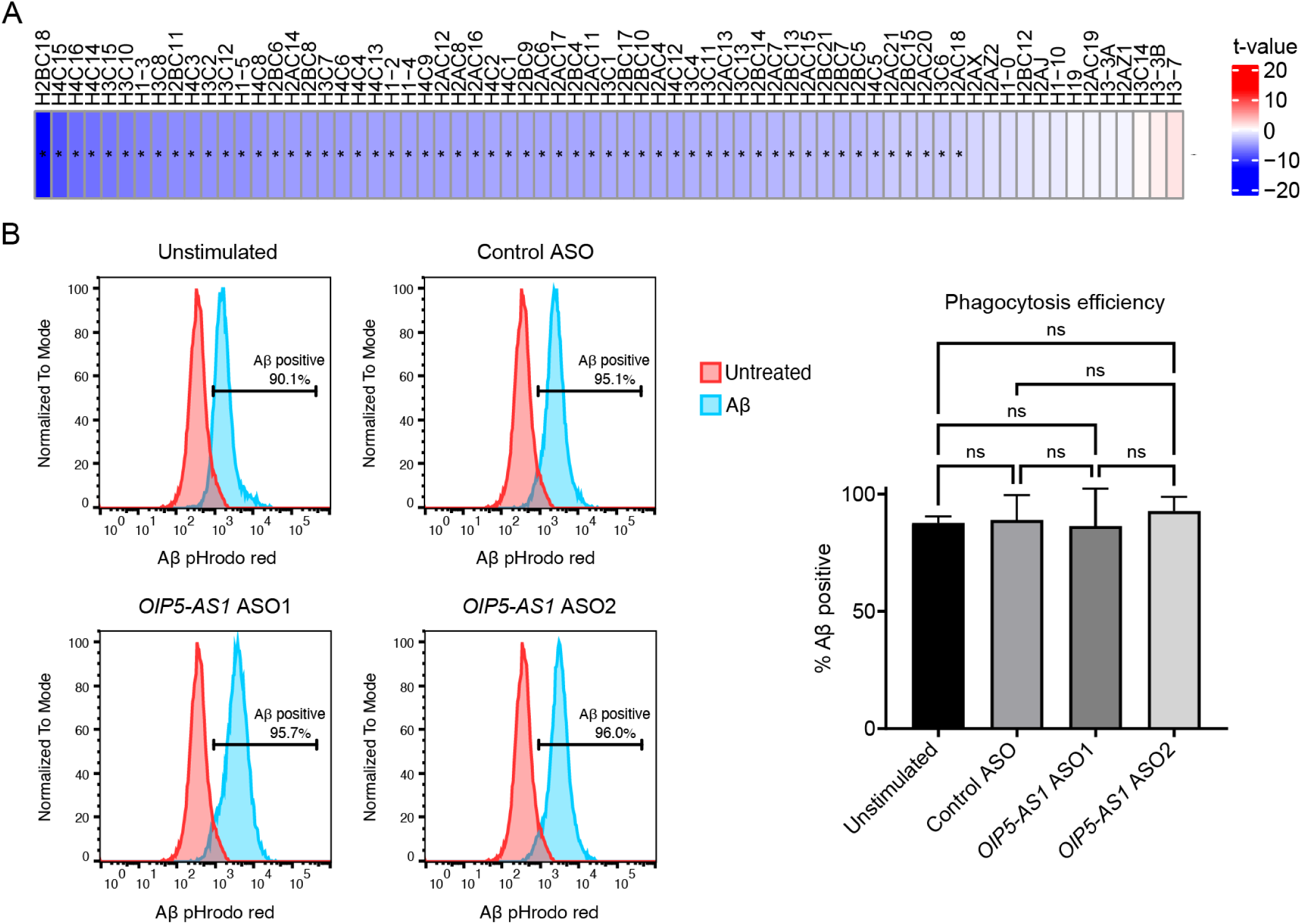
*OIP5-AS1* knockdown results in widespread downregulation of histone genes and no apparent change in phagocytosis. **(A)** Heatmap of *OIP5-AS1*-KD vs. Ctrl t-values from RNA-seq. Fifty out of 69 detected histone genes are significantly downregulated (*p < 0.01). **(B)** Histograms of flow cytometry results following incubation of the indicated cells with pHrodo Red-labeled Aβ. The red peak indicates negative/background signal, and the blue peak indicates cells that have phagocytosed the labelled Aβ. The barplot on the right shows average percent uptake across 3 biological replicates, which was not significantly affected by *OIP5-AS1* knockdown.

## Supplementary Tables

Table S1. Subregion-and cell-type-enriched lncRNAs

Table S2. ST gene expression module assignments

Table S3. ST gene expression module preservation

Table S4. ST gene expression module FEA

Table S5. AD-differential lncRNAs (ST, bulk, snRNA)

Table S6. AD-differential GSEA of ST genes

Table S7. AD-differential gene sets including lncRNAs

Table S8. DEGs and GSEA of OIP5-AS1 KD cells

